# Microwave assisted acidification for green hydrogen and organic acids production

**DOI:** 10.1101/2023.11.18.567651

**Authors:** Maximilian Barth, Magdalena Werner, Pascal Otto, Benjamin Richwien, Samira Bahramsari, Maximilian Krause, Benjamin Schwan, Christian Abendroth

## Abstract

**Background:** The integration of anaerobic digestion into bio-based industries can create synergies, which help to make anaerobic digestion self-sustaining. Two-stage digesters with separate acidification stages allow to produce green hydrogen and short-chain fatty acids, which are promising industrial products. Heat shocks can be used to foster the production of these products. However, the practical applicability is oft not addressed sufficiently. The here presented work aims to close this gap.

**Methods:** Batch experiments were conducted in 5 litre double-walled tank reactors incubated at 37 °C. Short microwave heat shocks of 25 min duration and exposure times of 5 - 10 min at 80 °C were performed and compared to oven heat shocks. Pairwise experimental group differences for gas production and chemical parameters were determined using ANOVA and post-hoc tests. High-throughput 16S rRNA gene amplicon sequencing was performed to analyse taxonomic profiles.

**Results:** Heat shocking the entire seed sludge, the highest hydrogen productivity was observed at a substrate load of 50 g/l with 1.09 mol H_2_/mol hexose. With 1.01 mol H_2_/mol hexose, microwave assisted treatment was not significantly different from oven-based treatments. The study emphasised the better repeatability of heat shocks with microwave-assisted experiments, showing low variation coefficients averaging 29 %. Microwave pre-treatment indicates a high predictability and a stronger microbiome shift to Clostridia than the oven. The pre-treatment of heat shocks supported the formation of butyric acid up to 10.8 g/l in average and a peak of 24.01 g/l at a butyric/acetic acid ratio of 2.0.

**Conclusion:** Results show the suitability to heat shock the entire seed sludge rather than just a small inoculum, which makes the process more relevant for industrial application. A microwave-based treatment has proven to be a promising alternative to oven-based treatments, which ultimately might facilitate the implementation into industrial systems. The approach becomes economically sustainable with high-temperature heat pumps with a coefficient of performance (COP) of 4.3.

## 1 Background

Anaerobic digestion (AD) is a technology that enables the production of methane from various organic resources and residues, such as straw and corn stover [1], chicken manure [2], food waste [3]. The potential for generating methane from biodegradable waste presents an interesting prospect. However, it faces significant challenges when compared to established technologies that utilize fossil fuels. In Germany, the monetary shortcoming is counterbalanced based on the German Renewable Energy Sources Act [4]. Therefore, technological and conceptual advancements are required to make AD a self-sustained technology. A possibility to achieve this is to intertwine AD with existing bio-based industries creating synergetic effects. For instance, Sawatdeenarunat et al. discussed the possibility of producing valuable side products from extracted solids, the digestate or the biogas itself, as syngas, methanol, butanol and others [5]. A particularly promising approach to produce valuable side products is the application of two-stage digesters with a separated acidification stage. Using high loading rates, acidogenic microorganisms allow for the accumulation of short-chain fatty acids (C2 – C5), which are important platform chemicals [6]. Under specific conditions even a chain elongation can be achieved during the digestion process, yielding medium-chain carboxylic acids [7]. Another promising metabolite formed during AD is hydrogen [8]. Particularly, a two-stage digestion process is well suited for microbial hydrogen production [9]. Using acidic pre-treatment stages of anaerobic digesters for hydrogen production is a process, which is very similar to dark fermentation. Dark fermentation approaches are extensively described in the literature and mixed culture fermentation was applied for microbial hydrogen production [10–12]. AD involves around 300 operational taxonomic units, which cover 80 % of all reads from 16S rRNA gene amplicon high throughput sequencing of samples belonging to 32 full-scale digester plants [13]. Such a high diversity makes it difficult to achieve a stable microbiome that yields a predictable outcome of hydrogen. Another problem that occurs when coupling a dark fermentation stage with a methanogenic reactor is the immigration of methanogens. Although high organic loading rates (OLR) and, low pH levels tend to suppress methanogenesis [14, 15], the formation of methane during dark fermentation has been described before [16]. Interestingly, heat shocks can help to optimise this problem. Although methanogenesis is not fully suppressed, it has been indicated recently that heat shocks can reduce the amount of methane, which is produced in separated acidification stages [17]. The fact that heat shocks can simultaneously suppress methanogenesis and contribute to an enrichment of hydrogen-forming bacteria makes them particularly interesting. Although several research articles already address the application of inoculum heat-shocks to improve hydrogen formation, the authors of the present work have barely detected scientific articles, where the whole digesting mix was heat shocked. In respect to an industrial application, the treatment of the complete seed sludge would be more practical. The presented work closes this research gap by investigating the industrial suitability of heat-shocks for separated acidification stages heating up the entire seed sludge.

The utilisation of existing waste streams in the form of various fermentation residues from biogas production offers new cascading recycling stages through heat shock pre-treatment. For the first time, the application of microwaves was assessed in comparison to heat shocks in the oven. Further, the effect of heat shocks on the underlying microbiome and the co-production of volatile fatty acids in the acidification stage were investigated.

## 2 Methods

### 2.1 Experimental set-up

Six identical double-walled tank reactors (3.3 l working volume) from the company Lehmann-UMT GmbH (Germany) were applied (Figure 1A). The batch experiments were fed with different loading rates of 50, 75 and 100 g/l of sucrose. All reactors were operated at 37 ± 1 °C.

**Figure 1:**
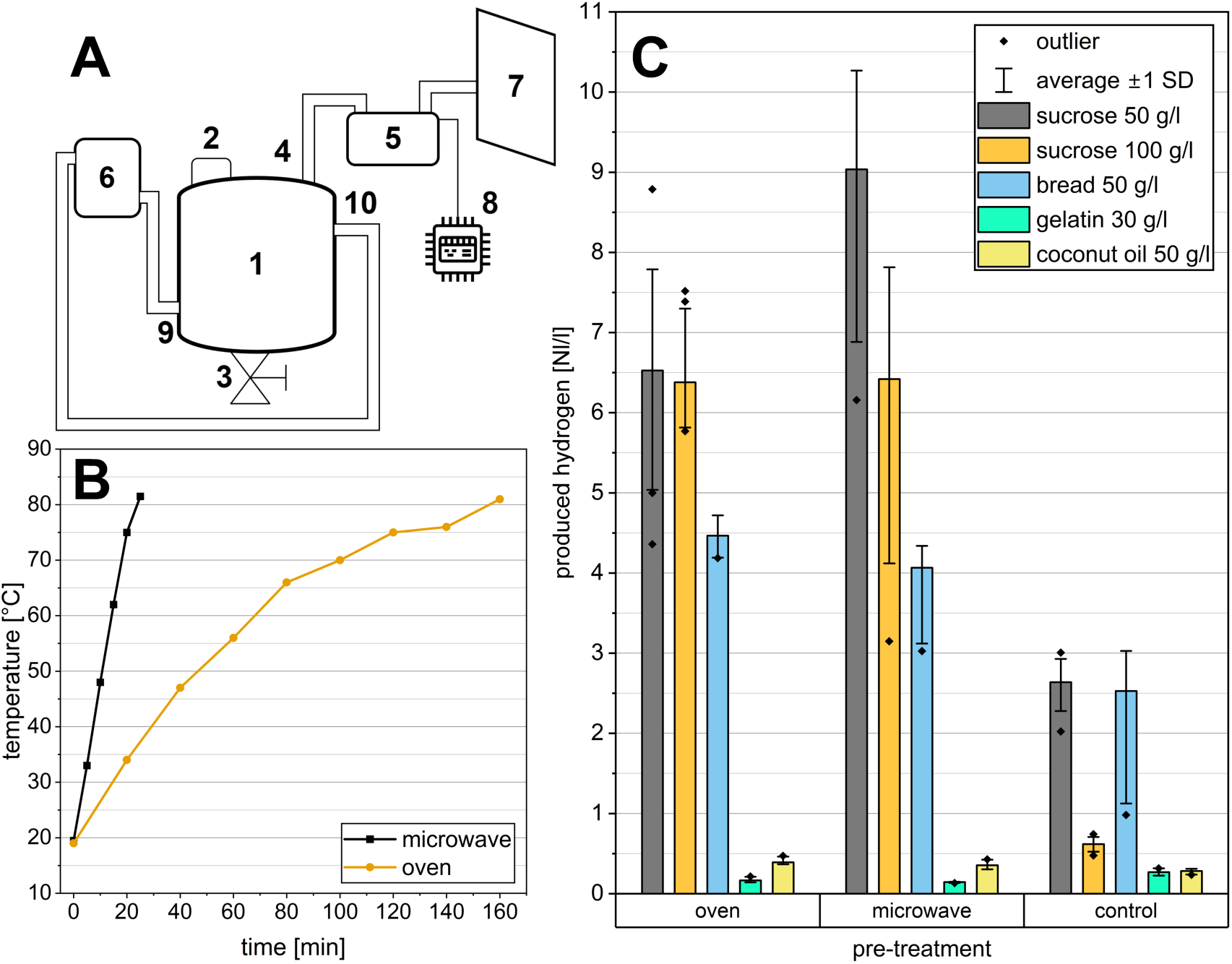
**(A)** Experimental set-up: double-walled reactors (1) with a volume of 5 l each. Each reactor was filled through an input port (2) and acidified hydrolysate was removed through a port at the bottom (3). For heating, a water pipe was connected to the input (9) and output (10) ports of the double-wall clearance. Water was heated using a thermostat (6). Produced gas was measured by a MilliGascounter (5), connected with a tube (4), and stored in a gas bag (7). The time-resolved gas production was recorded with an Arduino (8) connected to the gas counter. **(B)** Average incubation time for heat shocking up to a minimum of 80 °C for pre-treatment with oven (set to 130 °C) and microwave (set to 1000 W). **(C)** Substrate variation tests: Comparison of hydrogen production with sucrose, mixed rye bread, gelatine and coconut oil as feeding substrates and sewage sludge as inoculum. Both control and heat shock (oven and microwave) scenarios were tested.

The entire digestion mixture was subjected to heat shocks before being filled into the reactor. Different methods of heating were tested. For the heat shock, the slurry was heated using both oven and microwave. By oven, the sludge was incubated at 130°C in 1 litre Schott flasks for up to 3 hours. By microwave, the sludge was heated in 5 l measuring cups at 1000 W power output for only 25 min. Both methods were used to heat the sludge up to 80-90 °C, to eliminate hydrogen consuming microorganisms. The incubation times and heating curves are shown in Figure 1B. After the heat shock, the sludge was cooled down to 40 °C. Mohanakrishna and Pengadeth reported that heat shocks are often performed in the range of 65 to 121 °C and with exposure times between 1 to 10 h [18]. In the present work, the temperature of the heat shock is up to 80 °C ± 1 °C and the exposure times only at 5-10 min for the microwave and 10-20 min for the oven pre-treatment. These lower conditions were chosen based on the experiments of Wong et al. [19] with the aim to reduce the heat shock duration and exposure time even further. In their experiments, they had shown that effective heat shock pre-treatment of anaerobic sludge could be achieved at lower temperatures (65-85 °C) and for 45-60 min. Considering that the entire fermentation mix was heat treated in the present work, lower temperatures and shorter incubation times were preferred to be more energy efficient.

The experiments tested two conditions: feeding rate and heat shock method. Three feeding scenarios (50, 75, 100 g/l) were combined with three treatment methods (control, microwave, oven). To ensure high statistical confidence, each scenario was repeated at least six times, resulting in a total of 69 data points analysed. The data was composed of 29 control experiments, 20 oven pretreated experiments, and 20 microwave pretreated experiments.

### 2.2 Inoculum and Substrate

Dark fermentation experiments were performed, using digested sewage sludge as seed sludge. Substrates that are rich in proteins and fats were tested in a first set of experiments to evaluate their suitability in a heat shock context, in addition to carbohydrates (Figure 1C). Based on the better performance sucrose was chosen as the carbon source for the ongoing experiments. The sewage sludge was retrieved from a digester of a municipal wastewater treatment plant in Saxony, Germany. Table 1 shows the working parameters of the wastewater treatment plant.

**Table 1:**
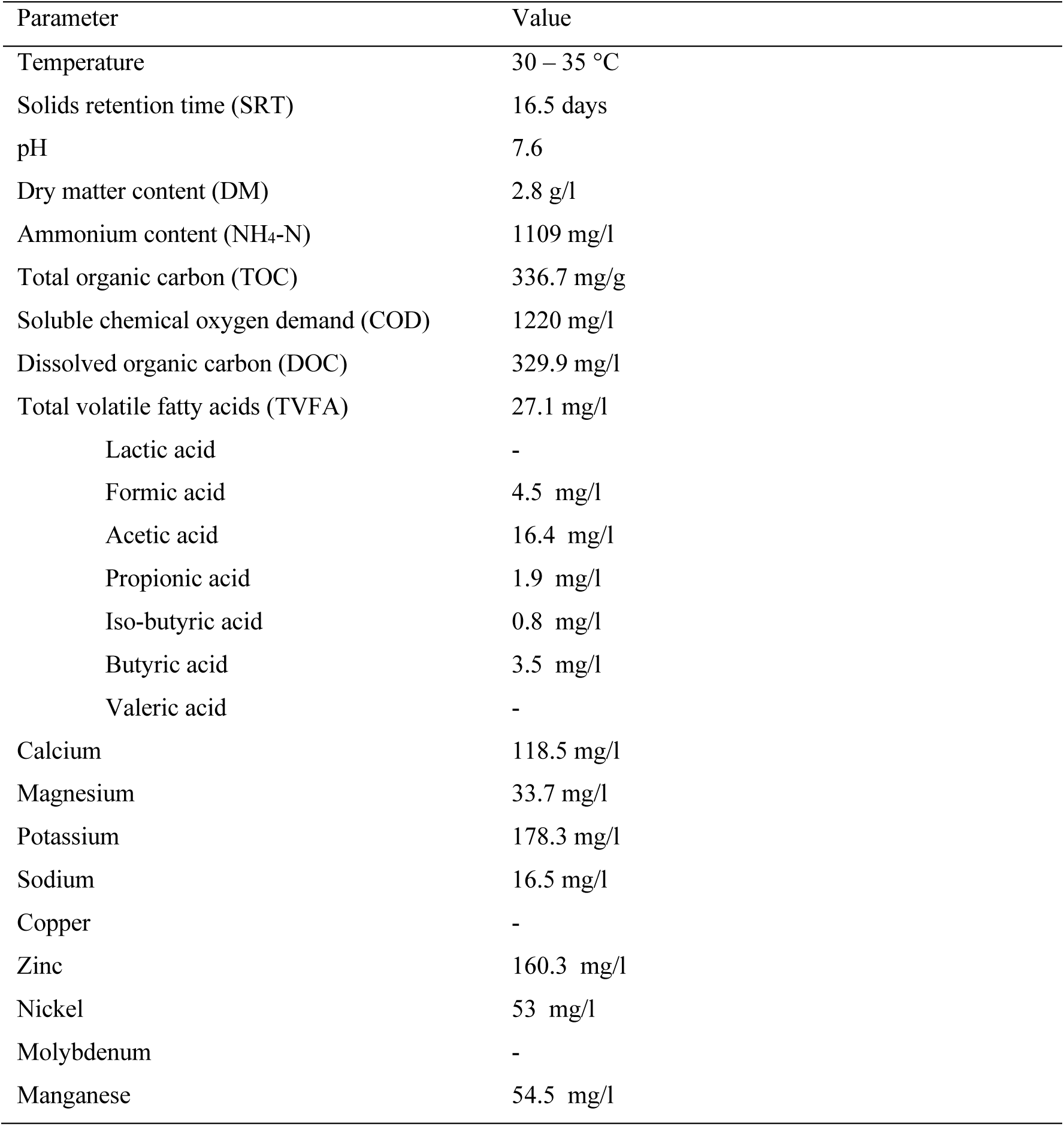
Working parameters and digested sludge parameters of the wastewater treatment plant in Saxony.

| Parameter | Value |
| --- | --- |
| Temperature | 30 – 35 °C |
| Solids retention time (SRT) | 16.5 days |
| pH | 7.6 |
| Dry matter content (DM) | 2.8 g/l |
| Ammonium content (NH<sub>4</sub>-N) | 1109 mg/l |
| Total organic carbon (TOC) | 336.7 mg/g |
| Soluble chemical oxygen demand (COD) | 1220 mg/l |
| Dissolved organic carbon (DOC) | 329.9 mg/l |
| Total volatile fatty acids (TVFA) | 27.1 mg/l |
| Lactic acid | - |
| Formic acid | 4.5 mg/l |
| Acetic acid | 16.4 mg/l |
| Propionic acid | 1.9 mg/l |
| Iso-butyric acid | 0.8 mg/l |
| Butyric acid | 3.5 mg/l |
| Valeric acid | - |
| Calcium | 118.5 mg/l |
| Magnesium | 33.7 mg/l |
| Potassium | 178.3 mg/l |
| Sodium | 16.5 mg/l |
| Copper | - |
| Zinc | 160.3 mg/l |
| Nickel | 53 mg/l |
| Molybdenum | - |
| Manganese | 54.5 mg/l |

### 2.3 Analytical methods

#### 2.3.1 Gas volume and composition

During the fermentation process, the produced gas was collected in gas bags (Tesseraux, Germany) and continuously quantified using a RITTER MilliGascounter (Bochum, Germany). Based on the guideline VDI 4630 from the Association of German Engineers (2016), the gas volume was normalised to standard temperature (273 K) and standard pressure (1013 hPa).

The gas composition, consisting of hydrogen (H_2_), carbon dioxide (CO_2_), methane (CH_4_) was determined using a BlueVary gas analyser from BlueSens (Herten, Germany). To ensure anaerobic conditions, the oxygen (O_2_) and hydrogen sulphide (H_2_S) content was measured using the X-AM 8000 gas detector by Dräger (Lübeck, Germany).

The amount of hydrogen sulphide was very low (usually a few ppm), and therefore, it was neglected in the later results. To calculate the overall yield, the gas contained in the headspace was considered as well. The yield of hydrogen was estimated using equation 1 considering the density and the molar mass of hydrogen, the input mass and molar mass of the substrate (sucrose) and the number of hexose molecules per molecule of sucrose. The equation assumes that the substrate is completely consumed.

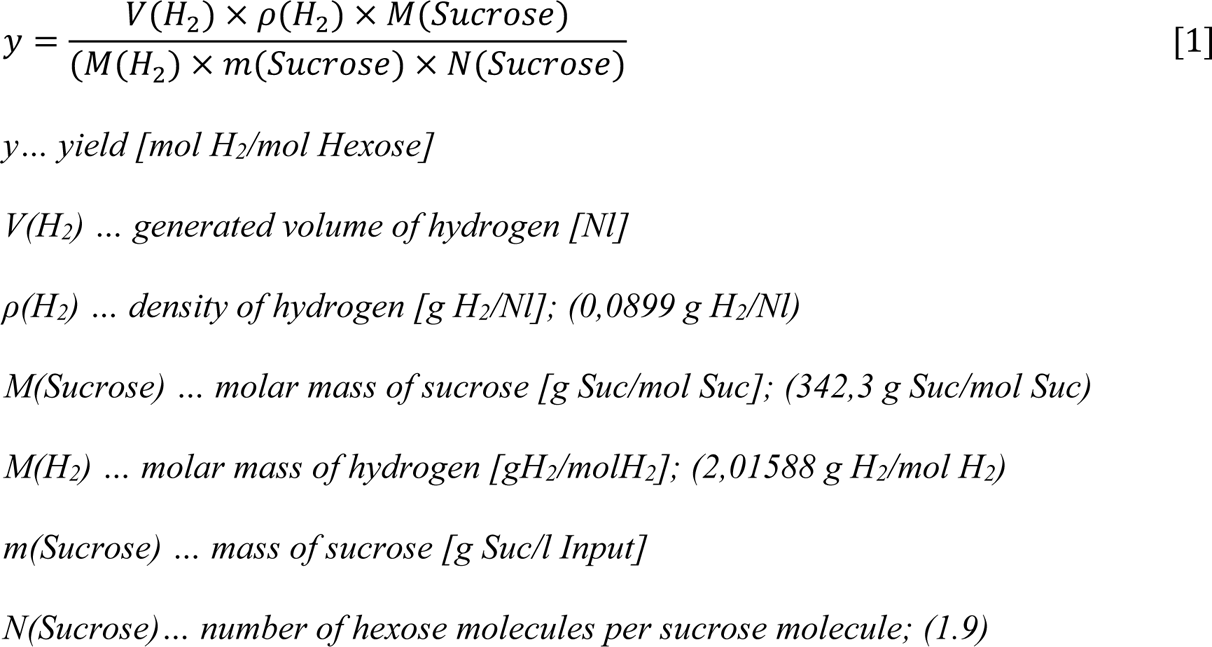

#### 2.3.2 Digestate analysis

For chemical analysis of the digestate, samples of 300 ml of the initial sludge were taken before heat shock and samples from each reactor after a complete fermentation phase. The solubilised COD was measured in the liquid phase after centrifugation at 1200 relative centrifugal force (RCF) and subsequent vacuum filtration through a 0.2 µm cellulose-acetate filter (Sartorius AG, Göttingen, Germany). The COD was determined using the Spectroquant COD kit (VWR, Germany) according to the manufacturer’s guidelines. The spectrum of VFAs including lactic acid, formic acid, acetic acid, propionic acid, iso-butyric acid, butyric acid and valeric acid was determined by ion chromatography (IC). A cation exchange column (Metrosep Organic Acids 250/7.8 column; Model: 882 Compact IC plus, Metrohm AG, Herisau, Switzerland) was used for this purpose. The mobile phase had a concentration of 0.6 mmol/l of perchloric acid and 10 mmol/l of lithium chloride. The detection limit was 0.25 mg/l. TVFAs were analysed as the sum of all detected VFAs. Initially, heavy metals were measured by microwave digestion and Inductively Coupled Plasma (ICP) (PerkinElmer Optical-Emission Spectrometer Optima 8000/ S10 Autosampler). Nitrogen contents including total Kjeldahl nitrogen (TKN) and NH_4_-N, were determined according to ISO 5663.

### 2.4 Metagenomics analysis of the microbial community

For microbiome analysis with 16S rRNA gene amplicon high-throughput sequencing, an aliquot of 3 ml of each sample stored in ethanol was centrifuged and washed with sterile phosphate-buffered saline (PBS) until the supernatant was clear. DNA was extracted from the resulting pellets using a Qiagen DNeasy PowerSoil Kit. DNA was quantified using the Qubit 1x dsDNA Kit from ThermoFisher. The extracted metagenomic DNA was used to amplify the hypervariable region V3-V4 of the 16S ribosomal RNA gene. The conserved regions V3 and V4 (470 bp) of the 16S rRNA gene were amplified using PCR cycling: initial denaturation at 95 °C for 3 min; 25 amplification cycles (20 s at 98 °C, 15 s at 60 °C, 30 s at 72 °C); and 5 min extension at 72 °C. The following primers were used:

- 341F (5’ ACACTCTTCCCTACACGACGCTCTTCCGATCT NNNNN CCTAYGGGRBGCASCAG 3’) and
- 806R (5’ GTGACTGGAGTTCAGACGTGTGCTCTTCCGATCT GGACTACNNGGGTATCTAAT 3’).

Amplification was performed using the Thermo Scientific Phusion High-Fidelity DNA polymerase kit. The purification of the amplicons was carried out after each PCR with the 1x v/v AmPure XP beads from Beckmann Coulter. The purified 16S amplicons were indexed with a second PCR containing TruSeq Illumina Indexing Primers (cycling conditions 98 °C 30 s denaturation, 98 °C 15sec, 65 °C 75 s, 10 cycles, final elongation 5 min 65 °C, 4 °C overnight). Final libraries were again purified with 1x v/v AmPure XP beads and quality-assessed on an Agilent FragmentAnalyzer (1-6000bp NGS Kit). Resulting libraries were equimolarly pooled and sequenced. Sequencing was performed using 2X250pb or 2 × 300pb paired-end cycle runs on an Illumina MiSeq instrument. The Illumina raw sequences were loaded into Qiime2 (v. 2021.2.0) [20]. The quality of the sequences was checked with the Demux plugin and the DA-DA2 pipeline integrated in Qiime2. It was used to trim and join the sequences, remove chimaeras and detect amplicon sequence variants (ASVs) (> 99.9 % similarity). The taxonomy of each sequence variant was determined using SILVA [21] as the reference database for taxonomic assignment.

### 2.5 Statistical analysis

For the basic statistical evaluation of the test series, each with at least six measured values, the median was selected to better represent the tendency of the individual test results. To test the experimental groups for statistically significant group differences, the Levene test was first used as a test procedure for homogeneity of variance. Unlike the Bartlett test, this test is significantly more robust when populations are not normally distributed [22]. For experimental groups that did not have homogeneity of variance, a WELCH ANOVA was performed [22, 23]. The groups with homogeneity of variance were compared using classic ANOVA. Pairwise experimental group differences were determined using post-hoc tests. For this purpose, the Tukey-Kramer test [24, 25] and, for groups without homogeneity of variance, the Games Howell test was used [26]. For all tests, the significance level was set at α = 0.05.

## 3 Results

### 3.1 Dark mixed culture fermentation with different substrates

The hydrogen production for the fatty substrates resulted in general low rates with maximums of 0.32 Nl/l for gelatine and 0.47 Nl/l for coconut oil. The protein-rich mixed rye bread produced a maximum hydrogen yield of 4.46 Nl/l with oven pre-treatment and 4.07 Nl/l with microwave pre-treatment. However, it was below the results obtained with 50 g/l and 100 g/l sucrose. The respective hydrogen yields are shown in Figure 1C. In the control group without treatment, a hydrogen yield of only 0.62 Nl/l was achieved when loaded with 100 g/l sucrose. In contrast, both heat shock scenarios resulted in a hydrogen production of 6.42 Nl/l at this high feeding overload. Highest hydrogen production was achieved with 50 g/l sucrose and oven heat pre-treatment of the sludge. Overall, the heat-treated trials produced twice as much total gas compared to the untreated control.

### 3.2 Combination of heat shock pre-treatment and high feeding rates

As shown in Figure 1C, fats and proteins could not match the hydrogen production rates of sugar in the dark fermentation combined with heat shock. Therefore, subsequent experiments were focused on very high feeding rates of sucrose as substrate. As shown in Figure 2, different sucrose loading rates (50, 75 and 100 g/l) were compared in terms of hydrogen production, to determine the optimal substrate concentration. Assuming an idealised uniform consumption of the substrate, this would correspond to loading rates of 7, 11 and 14 g COD/l*d. The highest average hydrogen yields in mol H_2_ per mol hexose were obtained within the heat shock-treated experiments, with a loading of 50 g/l sucrose (Figure 2C). Microwave and oven pretreated sludges showed hydrogen formation efficiencies of 0.51 and 0.95 mol H_2_/mol hexose, respectively. The peak values for the hydrogen efficiencies were measured at 1.01 (microwave) and 1.09 mol H_2_/mol hexose (oven) both with a loading rate of 50 g/l sucrose.

**Figure 2:**
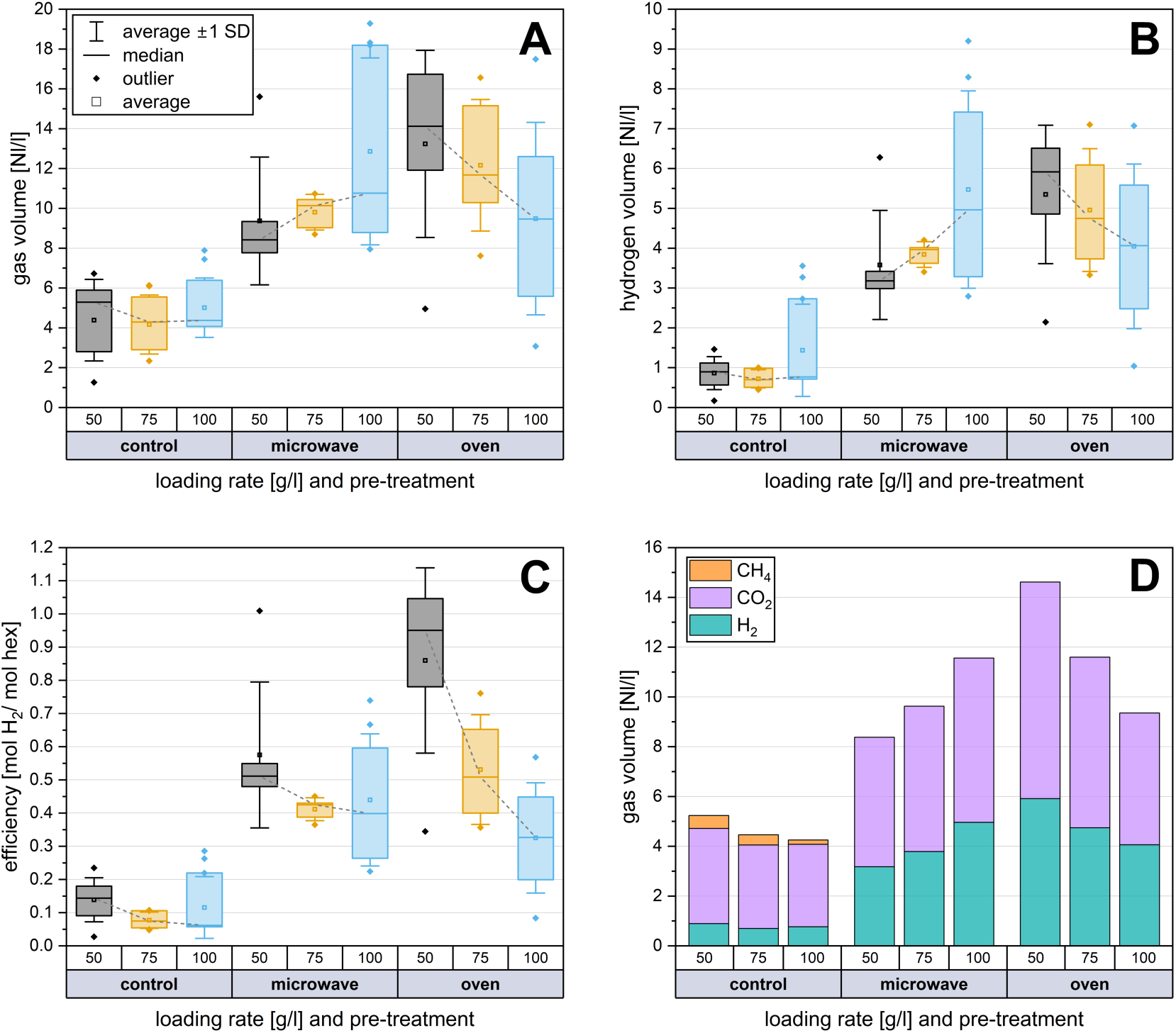
Application of heat shocks to optimise hydrogen productivity: Different concentrations of substrate (sucrose) and different pre-treatments (oven and microwave) were compared. **(A)** Maximal gas productivity and **(B)** the highest hydrogen production were reached with substrate concentration of 50 g/l and pretreated with oven heat shock. All fermentations were performed at least 5 times. **(C)** The highest hydrogen efficiency was achieved with 50 g/l sucrose both in microwave and oven experiments. **(D)** shows the gas composition depending on the feeding amount and pre-treatment. The median values were linked to a trend line in order to better visualise the effects of the different loading rates.

The untreated control showed the highest median hydrogen formation at 100 g/l sucrose with 0.2 mol H_2_/mol hexose, corresponding to 0.77 l hydrogen per litre working volume. In the heat-treated experiments, the lowest median hydrogen yield was produced with 100 g/l sucrose, with 0.39 and 0.33 mol H_2_/mol hexose for microwave and oven pre-treatment. As depicted in Figures 3A and B, increasing the feeding rates led to a rise in both the total volume and the absolute quantity of hydrogen, but only in the experiments that underwent microwave pre-treatment. Conversely, in the oven-treated experiments, the volumes decreased, leading to an inverse effect. Consequently, the efficiency of hydrogen formation in the oven experiments dropped significantly by approximately two-thirds from 50 g/l to 100 g/l. However, the decrease in hydrogen yield with microwave pre-treatment was only 28 %. In the untreated control experiments, there was only a slight change in gas volumes.

**Figure 3:**
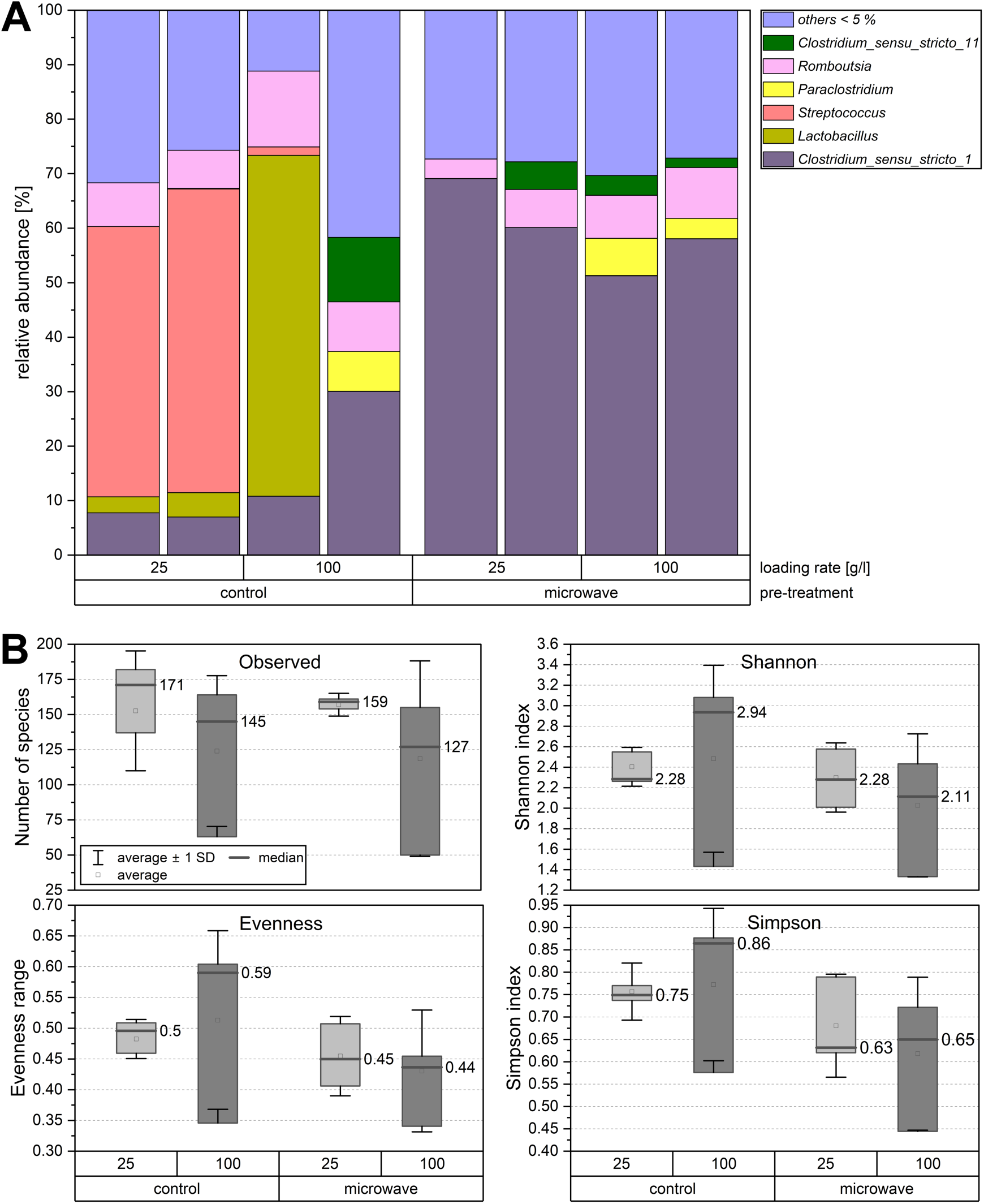
**(A)** Microbial community on genus level as relative abundances differentiated according to pre-treatment and sucrose loading rate. Only the most abundant microorganisms with abundance > 5 % are presented. Others with abundances below 5 % are summarised as others < 5.0 %. **(B)** Alpha Diversity of the microbiome for genus level data of microwave and control samples, loading rates of 25 and 100 g/l of sucrose.

Figure 2A illustrates that the heat shocks significantly increased both the total gas volume and absolute hydrogen production (Figure 2B) and improved the hydrogen formation efficiency (Figure 2C) compared to the untreated control experiments. In the heat shock trials (both oven and microwave), methane production was successfully eliminated (Figure 2D).

As a result of the heat treatments, the relative standard deviation of the hydrogenation efficiency increased by 49 % on average in the microwave pre-treatment and by 70 % in the oven pre-treatment. The standard deviation of the oven tests was on average 32 % higher than in the microwave tests. The pre-treatment with the oven was overall related to higher gas production compared to the microwave. However, the boxplots of the series of experiments show that the microwave shocks reduced the scatter of the results, an effect that was not observed in the experiments with oven treatment.

Distinct observations were made at a loading rate of 50 g/l sucrose. Here, the oven demonstrated a significant increase in absolute hydrogen production and consequently in hydrogen formation efficiency (Figure 2B and 2C). A median of 5.92 l H_2_/l sludge was produced, corresponding to a hydrogen yield of 0.95 mol H_2_/mol hexose. For both heat shock methods, the median hydrogen formation efficiency shows a decreasing trend as the loading rate increases. This pattern is reflected in the total volume and hydrogen volume for the oven-based heat shock experiments. In contrast, specific hydrogen production increases with increasing feeding rate in the microwave pre-treatment experiments (Figure 2C). Concurrently, the microwave-assisted hydrogen production efficiency remains more constant, decreasing by only 22 % from 50 g/l to 100 g/l (65 % decrease for oven pre-treatment). The distribution of gas volume indicates that in all experiments involving heat shock pre-treatment, the total hydrogen content in the gas increased to approximately 40 % for microwave and 41 % for oven pretreated. This is a significant increase when compared to the median of the control at 18 % H_2_ (Figure 2D). While in microwave-treated approaches the relative hydrogen fraction increased slightly from 38 % to 42 %, it remained constant at 40 % to 42 % in the oven-treated experiments. In both heat shock methods, the highest relative hydrogen content was obtained in the experiments with 100 g/l sucrose. Methane was not produced in any of the heat-treated trials. The residual portion of the gas mixture consisted entirely of carbon dioxide.

In general, thermal pre-treatment increased the hydrogen production capacity of anaerobic sludge in dark fermentation experiments by at least a factor of 3.6 for microwave and a factor of 5.3 for oven compared to untreated experiments. The maximum increase of 6.8 times was obtained in oven-treated trials at 75 g/l loading rate and in microwave-assisted trials by 6.5 times at 100 g/l loading rate (microwave: 50 g/l - 3.6 times, 75 g/l - 5.4 times; oven: 50 g/l - 6.6 times, 100 g/l - 5.3 times). The strongest increase of 1.9-fold in the oven-treated trials over those with microwave treatment was achieved at a loading rate of 50 g/l. However, this did not result in statistically significant differences between microwave and oven pre-treatment in terms of overall hydrogen volume (p-values for all loading rates in Figure 4A). At 100 g/l feeding rate, no significant group differences between the control and oven for total gas volume (p = 0.08) and hydrogen content (p = 0.14) could be determined. For both the 50 g/l and 75 g/l loading rate experiments, there were statistically significant differences (p < 0.05) between both heath shock scenarios and control for the parameters hydrogen volume, hydrogen fraction, efficiency, and total gas volume.

**Figure 4:**
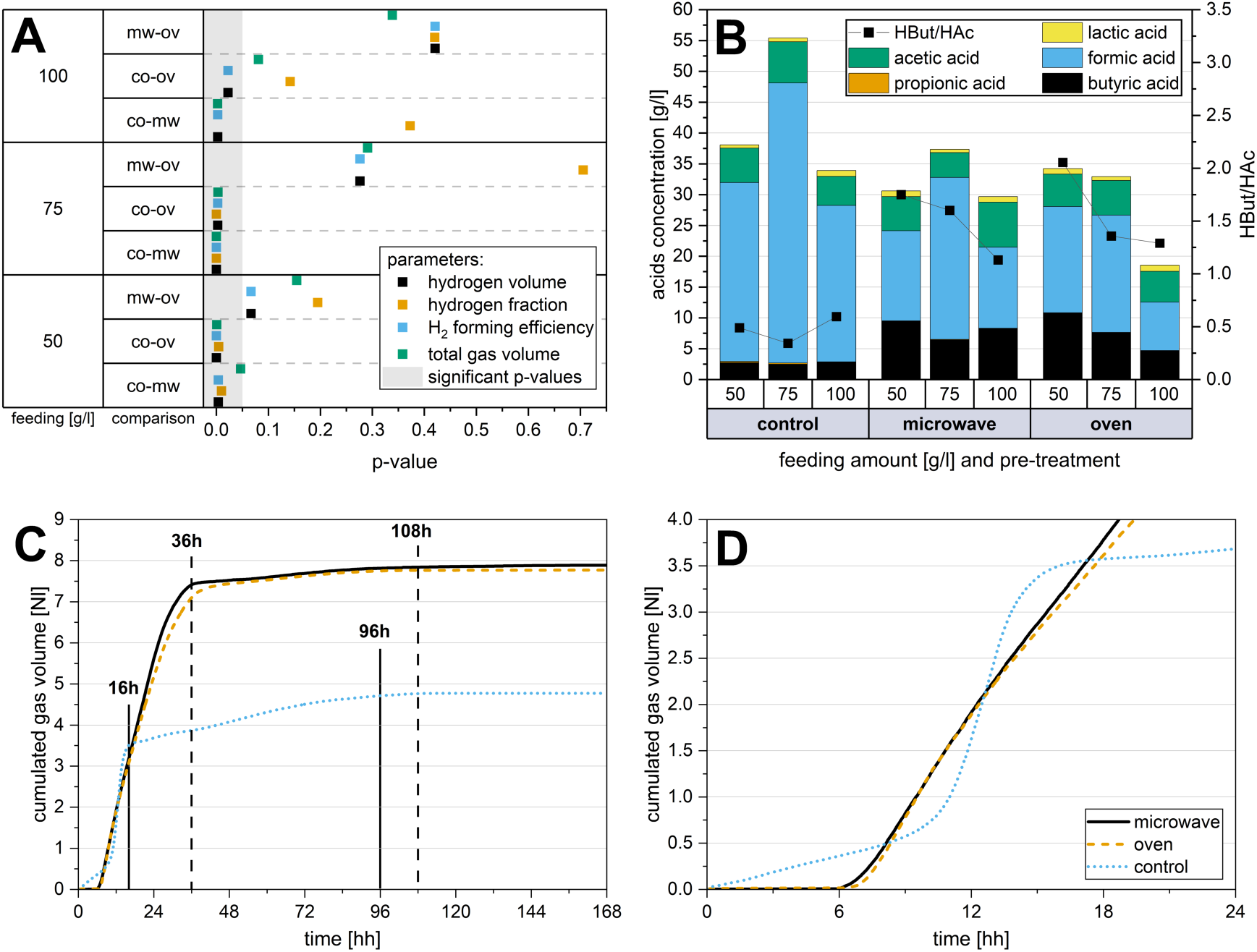
**(A)** Graphical representation of statistical significance (significant for p< 0.05) of group comparisons for hydrogen and total gas volumes, hydrogen formation efficiency and hydrogen fraction. The marker in the grey area indicates a significant difference between the respective comparison groups. **(B)** Production of volatile fatty acids (median) differentiated by treatment method and feeding amount. Volatile fatty acids were analysed in triplicates in all cases. **(C)** Exemplary course of gas production over seven days and **(D)** first 24 hours for untreated (control) and heat shock treated (oven and microwave). Performed with sewage sludge and 75 g/l sucrose as feeding substrate.

### 3.3 Microbiome taxonomy

The effect of microwave heat shock on the taxonomic profile and diversity of the microbiome was determined using 16S rRNA gene amplicon high-throughput sequencing. To show the range of microbiome changes, the edges of the test spectrum were defined and analysed at 25 g/l and 100 g/l sucrose. The gas formation efficiency, composition, and acid formation with 25 g/l sucrose were generally similar to the results obtained with 50 g/l but were slightly lower overall. The purpose of evaluating the microbiome at loading rate of 25 g/l was to reflect the taxonomic shifts as a result of overfeeding less strongly. Microwave treatment effectively eliminated non-spore formers and homogenised the microbiome towards the class Clostridia. In experiments with 100 g/l overfeeding, a similar effect was observed, but with slightly lower abundances of Clostridia. With a loading rate of 25 g/l sucrose Clostridia predominantly colonised the microwave-treated substrates at an average abundance of 91 % (100 g/l – 84 %). As depicted in Figure 3A, the microbial growth community in the microwave-treated reactions was composed of 51 % to 69 % *Clostridium sensu stricto 1*. In contrast, the genera *Streptococcus* and *Lactobacillus* dominated the control experiments, making up 49 % to 56 % and 63 % of the total microbiome. The heat shock succeeded in completely suppressing *Lactobacillus*, a lactic acid producer. Larger amounts of the genus *Romboutsia* were found in all scenarios with an average abundance of 8 %. The genus *Paraclostridium* was found in highly overfeeded experiments only with abundances of 4 % to 7 %. Microwave heat shock pre-treatment resulted in a reduction of observed species diversity by an average of 15 % (25 g/l) and 20% (100 g/l) compared to the control, as shown in Figure 3B. While the Shannon index showed no difference in diversity for 25 g/l, the Simpson index indicated a reduction in species diversity for both loading rates in the microwave-treated trials. Control scenarios exhibited a higher evenness, while microwave treated microbiomes pointing to a highly uneven distribution of species. The strong shift towards *Clostridium sensu stricto 1* and the decrease in absolute species number in the heat-treated experiments led to a more uneven distribution of species. This generally low equal distribution is also reflected in the taxonomic profile, which shows that an average of 28 % of the microbiome consists of smaller groups of microorganisms with less than 5 % relative abundance (Figure 3A).

### 3.4 Production of volatile fatty acids

In general, the TVFA production was approximately 30 to 40 g/l for heat-shocked experiments and between 35 and 55 g/l for non-pretreated ones as a result of the higher formic acid production. The thermal pretreated experiments demonstrated a significant shift of the produced acids towards butyric acid. In heat-shocked experiments, the molar ratio of butyric acid to acetic acid (HBut/HAc) was in a range from 1.13 to 2.05 (Figure 4B). Without thermal pre-treatment, the HBut/HAc ratio was between 0.34 and 0.59. At a loading rate of 100 g/l of sucrose and due to microwave pre-treatment, the butyric acid concentration increased 2.95-fold up to 8.29 g/l. In contrast, the concentration of formic acid decreased to 13.14 g/l (1.93-fold). The heat shock pre-treatment reduced the accumulation of formic acid over all loading rates by 1.85-fold average. With the oven pre-treatment and 50 g/l loading rate, the highest average butyric acid concentration was reached at 10.80 g/l followed by microwave with 9.48 g/l. Microwave pre-treatment led to a single peak concentration of 24.01 g/l butyric acid at a loading rate of 50 g/l sucrose. The amount of acetic acid was hardly influenced by the pre-treatment, neither by the oven nor the microwave. Of the seven VFA analysed, formic, acetic and butyric acid were the most frequent represented (in total >95 %). In the untreated controls, 2.73 g/l butyric acid and 29.03 g/l formic acid were formed at the same loading rate of 50 g/l. None of the experiments contained larger amounts of propionic (maximum of 0.18 g/l) or lactic acid (maximum of 1 g/l). Significant concentrations of iso-butyric acid (0.02 g/l) and valeric acid (0.01 g/l) could not be determined in any of the tests. After an experimental period of seven days, pH values of 3.98 on average were measured in all experiments. At the start of the experiments, the pH value was 7.56.

### 3.5 The kinetics of hydrogen production after heat shock application

In the untreated experiments, gas production starts immediately after adding the substrate (Figure 4C and 4D, with 50 g/l sucrose as substrate). This was followed by a rapid accumulation of biogas over the next 7 hours. After approximately 16 hours, a noticeable plateau in the gas production was reached, which comes to an almost complete standstill after about 96 hours. In contrast, total gas production in the heat shock experiments did not start until 6 hours after feeding but then ascends steeply and continuously until the 36th hour. The subsequent rise is significant less steep compared to the control experiment and nearly ceases entirely after 108 hours. This pattern was observed with both pre-treatment, oven and microwave. The rate of gas formation in the heat-treated trials was less steep compared to the untreated control (Figure 4C), yet the robust gas formation remained active for almost twice as long (30 hours) compared to the control trials (16 hours). The experiments demonstrated that heat shock pre-treatment leads to more consistent and prolonged gas production, significantly increases the total gas volume, albeit with a delay in gas production for the initial six hours.

The statistical significance was assessed on the basis of an ANOVA (Figure 4A) in order to better recognise whether the differences between the various treatments were significant. For example, at a loading rate of 100 g/l of sucrose, there is no significant difference between microwave and oven assisted pre-treatment regarding hydrogen volume, total gas volume, efficiency and hydrogen percentage (Figure 4A).

## 4 Discussion

### 4.1 Influence of substrate and feeding rate

Several research papers conclude that carbohydrate-rich substrates such as sugar, starch, or cellulose are especially effective for dark fermentation [27–29]. In addition to testing these suitable substrates, protein-rich and fat-containing substrates were also tested. The aim was to determine the suitability or unsuitability of these substrates in combination with heat shock pre-treatment of the entire initial sludge. The microbiome generated by the heat shock pre-treatment was not able to effectively metabolise the alternative substrates.

### 4.2 Application of shorter heat shocks to high-loaded dark fermentation processes

Saady [30] states that inhibiting H_2_-consuming microorganisms, such as hydrogenotrophic methanogens, homoacetogens, lactic acid bacteria, propionate-producing bacteria, and sulphate reducers, is a crucial step in H_2_ production by dark fermentation in mixed microbial communities. In many studies, heat shocks are used to generate a small volume of enriched hydrogen-producing microbiome, which is then inoculated into a medium [31–33].

The study carried out here has shown that the use of already fermented sewage sludge from anaerobic biogas production expands the usability of this waste stream in terms of cascade recycling and increasing its industrial relevance. This strategy allows the selection of spore-forming hydrogen-producing microorganisms and the inhibition of non-spore-forming hydrogen microorganisms from heterogeneous microflora. Processes involving heterogeneous microflora are considered more suitable than pure cultures due to their simpler process control and efficient substrate conversion, provided that the medium undergoes pre-treatment [34]. Pineda-Muñoz et al. succeeded in generating high hydrogen yields by combining heat shock and ultrasonic treatment but did not completely suppress methane production despite nearly identical heat shock conditions [35]. Singhal and Singh investigated the effect of different microwave intensities on an inoculum [33]. In contrast to the study conducted here, they did not treat the entire sludge, but only a small inoculum of 20% of the total volume. Their focus was on the different levels of irradiation and not on the heat shock itself, which is why they carried out the treatment for only 5 minutes. They achieved a maximum hydrogen yield of 14 mmol H2/mol sucrose with microwave pretreatment at power levels from 160 W to 800 W.

With the shorter microwave heat shocks (compared to conventional oven heat shocks) at a lower temperature, it was possible to completely inhibit methane production in the current experiments. There are other researchers, who also tried to inhibit methanogenesis due to heat shocks. For instance, Hasyim et al. conducted a shock at 105 °C for 20 min in an autoclave [36], while Wang et al. performed a 70 °C heat shock for 60 min with the support of free ammonia [32] to fully inhibit methane production. Although [36] and [32] were successful in inhibiting methanogens, the microwave has the advantage of heating the sludge medium 6.4 times faster than with an oven. The experiments conducted here have demonstrated that even shorter and less energy-intensive heat shocks can completely eliminate methane production and, thereby, facilitate biological hydrogen production from sewage sludge. It should be noted that the experiments were performed in batch. In a continuous set-up, it would still be conceivable that methanogens would gradually adapt to the temperature shocks.

The recorded hydrogen formation rates are relatively low compared to more recent studies [37–39], but it is important to note that the present results were achieved despite heavy overfeeding. It must be considered that the current values correspond to the median of a minimum of six tests each. Compared to other works, the present results stand out due to the high number of replicates, which enabled a more robust and meaningful statistical analysis to evaluate the generated data. Considering the increased formation of butyric acid (Figure 4B), a theoretical maximum hydrogen yield of 2 mol H_2_/mol hexose is possible [40]. With 1.09 and 1.01 mol H_2_/mol hexose in the present study, the achieved is much lower than the theoretical yield. Nevertheless, these results are in a similar range to Abdallah et al. who reached 1.1 mol H_2_/ mol hexose in a similar set-up. As in the here presented results, Abdallah et al. used no pH control, however, they used lower loading rates of 25 g/l [41]. Overall, the results indicate a high efficiency in the present experiments, even under high overloading rates and fast acidification due to missing pH control. In most studies, only triplicates were carried out and slightly higher hydrogen efficiencies were achieved. Compared to the high number of repeated experiments in the current study, other studies have a lower statistical certainty. Differences to other studies can arise due to the influence of the sludge used. It is possible that better results can be achieved with sewage sludge from other plants or with fermentation sludge from agricultural biogas plants. The composition of the sludge is often subject to seasonal fluctuations and therefore has a major influence on the fermentation performance.

Hydrogen yield could potentially be increased further by pH controlling, since pH values of 4.0 to 4.5 harm hydrogen production [42]. This could explain the decrease in hydrogen formation as the loading rate increases. Increased overfeeding acidifies the reactor much faster and inhibits hydrogen formation. Gioannis et al. investigated the effect of pH on hydrogen formation. They found out that the highest hydrogen production was attained at pH 6.0 [43].

Compared to the untreated control the hydrogen formation for 75 g/l loading rate was enhanced 6 times with microwave pre-treatment and 7.3 times with oven pre-treatment. These findings align with those reported by Baghchehsaraee et al. and O-Thong et al., who reported a 5.1 (80°C for 30 min) and 7.9-fold (100°C for 1 h) increase, as a result of thermal pre-treatments [42, 44]. The experiments carried out here showed a high standard deviation even if the experimental conditions stayed the same. However, the standard deviation of the hydrogen efficiency could be slightly reduced in trials with 50 g/l sucrose feed and strongly reduced with 75 g/l sucrose feed by using the microwave for pre-treatment.

### 4.3 Taxonomic shifts due to heat shocks

The stability of AD systems is related to a higher diversity of species [45]. However, by exposing the microbiome to extreme conditions, a robust microbiome can be created. In this study, stable hydrogen production was achieved despite a decline in the diversity of species recorded and a strong dominance of *Clostridium sensu stricto 1*. The process and the microbial diversity were limited by the high production of organic acids and the resulting rapid drop in pH. The predictability of the microbiome was significantly higher and more consistent due to the microwave heat shock pre-treatment. Additionally, the growth of *Clostridium sensu stricto 1* was more dominant than in the oven pretreated experiments. The strong shift in the microbiome towards Clostridia due to heat shock pre-treatment corresponds to the expectations according to the literature as they had higher relative abundances in thermophilic digesters and are known spore-formers [46]. Unfavourable conditions such as high temperatures make it difficult for non-spore-forming organisms to survive. As a result, almost exclusively the spore-forming Bacillota in the form of *Clostridiaceae* were enriched explosively in the heat-treated experiments. Tang et al. observed a similar shift to *Clostridium sensu stricto 1* in their experiments when the initial pH was increased from 4.0 to 11.0 [47]. In the present experiments the starting pH was at 7.5 and was rapidly decreasing to 4.0 to 4.5 after 48 h.

In the control experiments, the Bacilli were able to gain a growth advantage instead, as they were already more strongly represented in the initial sludge. Interestingly, the relative abundance of the class Bacteroidia remains unchanged in all experiments around 4 %. In comparative studies, a high relative abundance of Bacteroidetes is often found in single-stage biogas processes [48, 49]. In the experiments of Chen et al. [50] the relative abundance of Bacteroidetes increased strongly with organic overload. Bacteroidota-coupled secondary fermentation processes normally carry out biological acid degradation. Both the control and the heat shock experiments showed the same abundances of Bacteroidota, why it is likely that these were inhibited primarily by the overfeeding and less by the heat shock.

In the microwave pretreated experiments *Clostridium sensu stricto 1* showed a strong dominance. This genus describes strictly anaerobic fermenting spore-formers [51]. During the metabolisation of sugars and proteins, the main fermentation products are butyric and acetic acid [52]. The genus is also able to produce lactic acid and ethanol, propanol or butanol. In the present experiments, members of *Clostridium sensu stricto 1* were able to produce large amounts of butyric acid even under severe overfeeding rates, demonstrating their robustness under extreme conditions. They probably benefited more from the high sucrose load. The population of the genus *Romboutsia*, which comprises around 8% of the microbiome, remained almost unaffected by the heat shock pre-treatment with slightly higher abundances in the control experiments. According to Gerritsen et al. [53], *Romboutsia* is a potential acetogen, capable of assimilating carbon via the Wood-Ljungdahl pathway (reductive acetyl-CoA pathway). The consistent relative abundance of *Romboutsia* partly explains the high formic acid concentrations observed in all experiments (Figure 4B). They ferment sugar and metabolise it mainly to acetate, formate and lactate [54]. The control experiments showed that the microbiome was dominated by *Streptococci* at the lower loading rate of 25 g/l sucrose. These are facultative anaerobes that ferment glucose under anaerobic conditions to produce lactic acid, acetic acid, propionic acid, and formic acid [55]. Only the untreated experiments showed an increase in propionic acid concentrations compared to the initial concentrations in the inoculation sludge. It is probable that the propionic acid formers in the sludge, like *Streptococcus*, lack mechanisms to survive high temperatures, unlike spore formers. It is not reported that *Streptococci* produce any gas. This suggests that the carbon dioxide in the untreated experiments was produced by the other microorganisms present. The *Streptococci* growth was severely inhibited by heat shock pre-treatment. A small populations of the spore-forming *Paeniclostridium* [56] were only present in the heat-treated experiments. The results suggest that the class benefited from the heat shock treatment.

### 4.4 Bio-based production of short-chain organic acids

Large quantities of acids were formed in both the untreated and the heat shock-treated experiments. In combination with the low pH value at the end of the test series and the results of the microbiome analysis, it can be assumed that no effective acid degradation takes place in any of the tests and that instead there is a strong accumulation of acids. While the total acid concentration remains very similar regardless of the treatment and loading rate (except for the control with 75 g/l of sucrose), the proportion of individual acids changes as a result of the heat shock pre-treatment.

The butyric acid to acetic acid molar ratio curves show that the heat shock initialised the production of less acetic acid. Instead, the amount of butyric acid increased 2.7 times. The high HBut/HAc molar ratios for all heat shocked experiments (Figure 4B) indicate that hydrogen was produced via the butyrate pathway [35]. Without heat pre-treatment, the HBut/HAc ratio shows that hydrogen was accordingly formed via the acetate pathway.

While in experiments with microwave treatment the butyric acid concentration increases with heavy overfeeding (100 g/l), it decreases with increasing loading in experiments with oven pre-treatment. The heat pre-treatment reduced the variety of acids, as no propionic acid was formed anymore. Both lactic and propionic acid had very low concentrations, which corresponds to results of Fang and Liu [57]. Although, their formation is typical for processes with high loading rates [43, 58]. Therefore, it can be suggested that only a few propionic acid formers were already present in the seed sludge and that the conditions tested did not favour their growth. The experiments demonstrated the feasibility of producing high concentrations of bio-based butyric acid through dark fermentation processes by combining microwave heat shocks and heavy overfeeding. To increase the formation capacity of organic acids, it is recommended to improve buffering capacity, control pH value, and constantly remove organic acids. According to Pineda-Muñoz et al., decreasing pH leads to increased formation of acetic and formic acid [35]. Therefore, regulating pH could result in an even higher ratio of butyric acid.

### 4.5 Industrial relevance

The investigations carried out in this study indicates that one litre of heat-treated sludge can yield an average of 5.92 m^3^ of hydrogen per m^3^ sludge at a loading rate of 50 g/l. Subtracting the average values for the control from this, the heat treatment results in an additional gain of 5.02 m^3^ hydrogen per m^3^ of sludge. The thermal energy required to heat the seed sludge from 38 °C to 80 °C is 49.04 kWh/m^3^, considering that it takes 4.2 kJ to increase the temperature of 1 l sludge by 1 °C. The additional energy generated in the form of hydrogen due to the heat shocks corresponds to 54.18 kJ/l sludge, which corresponds to 15.06 kWh/m^3^ sludge. Based on the 49.04 kWh/m^3^ for the heat shock, the energy loss for the heat shock process is 33.98 kWh/m^3^. This amount of energy must at least be recovered using a heat pump, to make this concept energetically attractive.

Approved high-temperature heat pumps, which are currently available on the market, can achieve a coefficient of performance (COP) of 4.3 when heating to 80°C from a starting temperature of 20 °C (as per data from ENGIE Refrigeration GmbH in [59]). This implies that 1 kWh of electrical energy can produce up to 4.3 kWh of thermal energy. To generate 49.04 kWh of thermal energy, a heat shock requiring 11.4 kWh electrically would be needed, assuming a coefficient of performance (COP) of 4.3. At this COP, the heat shock generates an additional energy output of 3.66 kWh/m^3^ of sludge compared to the calorific energy of the additional hydrogen. The process would also be energetically profitable due to further energy losses during the heat exchange process with the sludge, which are not considered here. Future technological improvements that increase the COP could make enhanced hydrogen production through heat shocks even more sustainable.

In view of the different costs of hydrogen and electric energy, the economic appeal of the concept could maintain even if the hydrogen equivalent of one kWh is priced higher than electricity itself. Based on Machhammer et al. and Bukold, a good benchmark for the costs of hydrogen production is the average price for steam reforming of 1.70 €/kg H_2_ [60, 61]. The energy cost for a heat shock using a high-temperature heat pump with a COP of 4.3 would be 0.4 €/m^3^ of sludge, (based on an average industrial electricity price of 35 €/MWh). Selling the produced hydrogen at the same price of 1.70 €/kg would generate an average revenue of 0.91 € from the amount of hydrogen from one cubic meter of heat-shocked sludge. This corresponds to a profit of 0.51 €/m^3^ of sludge. Despite a net energy loss, the production of hydrogen through heat shocks proves to be more cost-effective compared to other alternative methods of hydrogen production. The current hydrogen costs of hydrogen production without steam reforming are quantified by Gerloff at 5.20 €/kg [62]. Lowering the heat shock temperature could significantly decrease the energy requirement and thus the essential costs, while potentially yielding similar or even higher hydrogen yields [44]. These calculations are based on ideal assumptions and the successful scaling of trials in an industrial setting, with a focus on electricity costs.

## 5 Conclusion

The presented study indicates that short heat shock pre-treatments and short exposure times of the entire seed sludge can trigger bio-hydrogen production from digested sewage sludge. The present study shows that short heat shock treatments and short exposure times of the entire bacterial sludge can trigger biohydrogen production from digested sewage sludge. Hydrogen production by dark fermentation using microwave heat shocks was statistically equivalent to oven heat shocks. Microwave pretreatment ensures high predictability of the microbiome. Despite a heterogeneous microbiome in the initial sludge, the heat pre-treatment led to a very homogeneous microbiome consisting of 83 % Clostridia. This specialised acidification microbiome was able to produce 1.01 mol H_2_/mol hexose and 24.01 g/l butyric acid even under high loading rates of 50 g/l sucrose. The high number of test repetitions indicates statistically significant results, which strengthens the relevance of these findings. By using already available high-temperature heat pumps and successful upscaling, heat shocks are energetically sustainable and economically competitive with electrolysis for hydrogen production. A significant by-product of microwave heat shocks was the production of significant amounts of butyric and formic acid. Through the further development of suitable extraction processes, short-chain organic acids based on petroleum could be replaced by bio-based acids from the further utilisation of digested sludge.

## 6 Declarations

### 6.1 Ethics Approval and Consent to Participate

Not applicable.

### 6.2 Consent for Publication

Not applicable.

### 6.3 Availability of Data and Materials

The genome data will be published in the gene bank database of the National Center for Biotechnology Information after acceptance of the paper. All other data generated or analysed during the trial are included in this published paper.

### 6.4 Competing Interests

The authors declare that they have no competing interests.

### 6.5 Funding

We are grateful for funding of the work by the German Ministry of Economic Affairs and Energy (grant numbers 49MF190057, 16KN070128 and 16KN070126).

### 6.6 Authors’ Contributions

*All authors contributed to the conception, design and methodology of the study. MB, MW, and SB performed the sampling and generated the chemical and operational data. MB, PO and MK carried out the 16s rRNA analysis, metataxonomics and multivariate analysis. MB wrote and revised the main manuscript and prepared the figures, and all authors revised earlier versions. All authors read and approved the final version of the manuscript*.

## 6.7 Acknowledgements

Not applicable.

## Supplementary information

**Figure S1:**
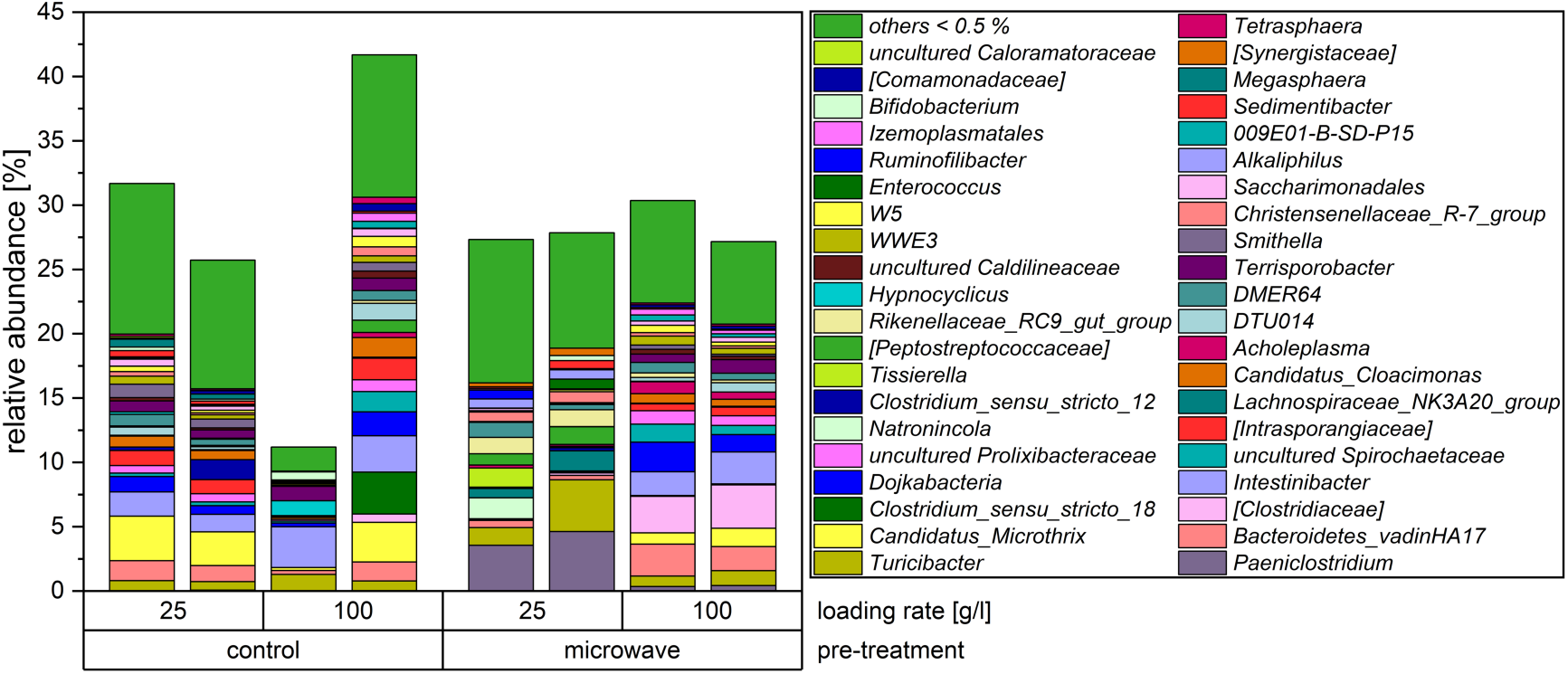
Microbial community on genus level as relative abundances differentiated according to pre-treatment (loading rates of 25 and 100 g/l). Only the microorganisms with abundance < 5 % are presented. Others with abundances below 0.5 % are summarised as others < 0.5 %.

**Figure S2:**
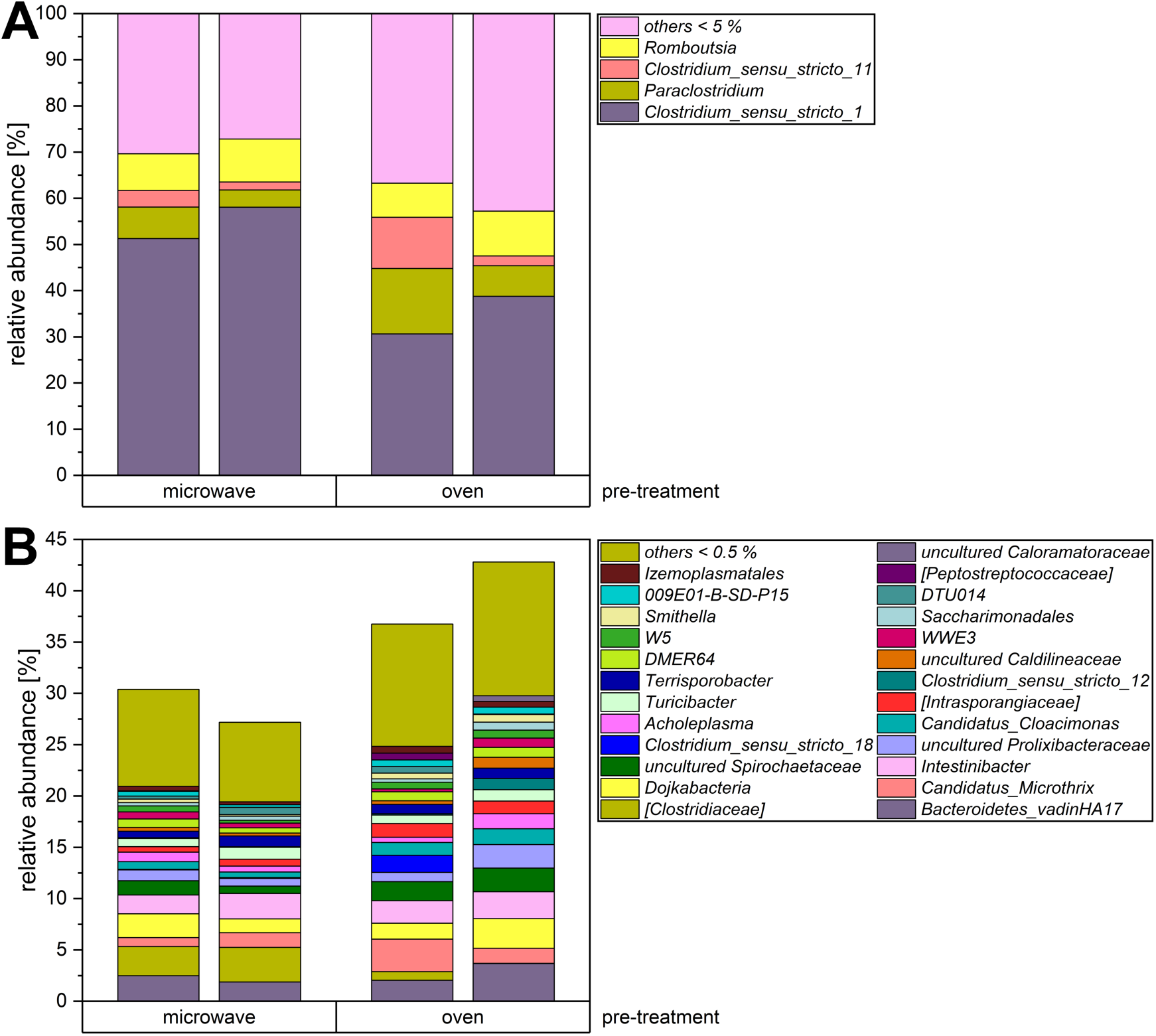
Microbial community on genus level as relative abundances differentiated according to pre-treatment. The loading rate was at 100 g/l sucrose. **(A)** Only the most abundant microorganisms with abundance > 5% are presented. Others with abundances below 5 % are summarised as others < 5.0 %. **(B)** Breakdown of minor microorganisms below 5 % abundance. Organisms with abundances below 0.5 % are summarised as others < 0.5 %.

**Figure S3:**
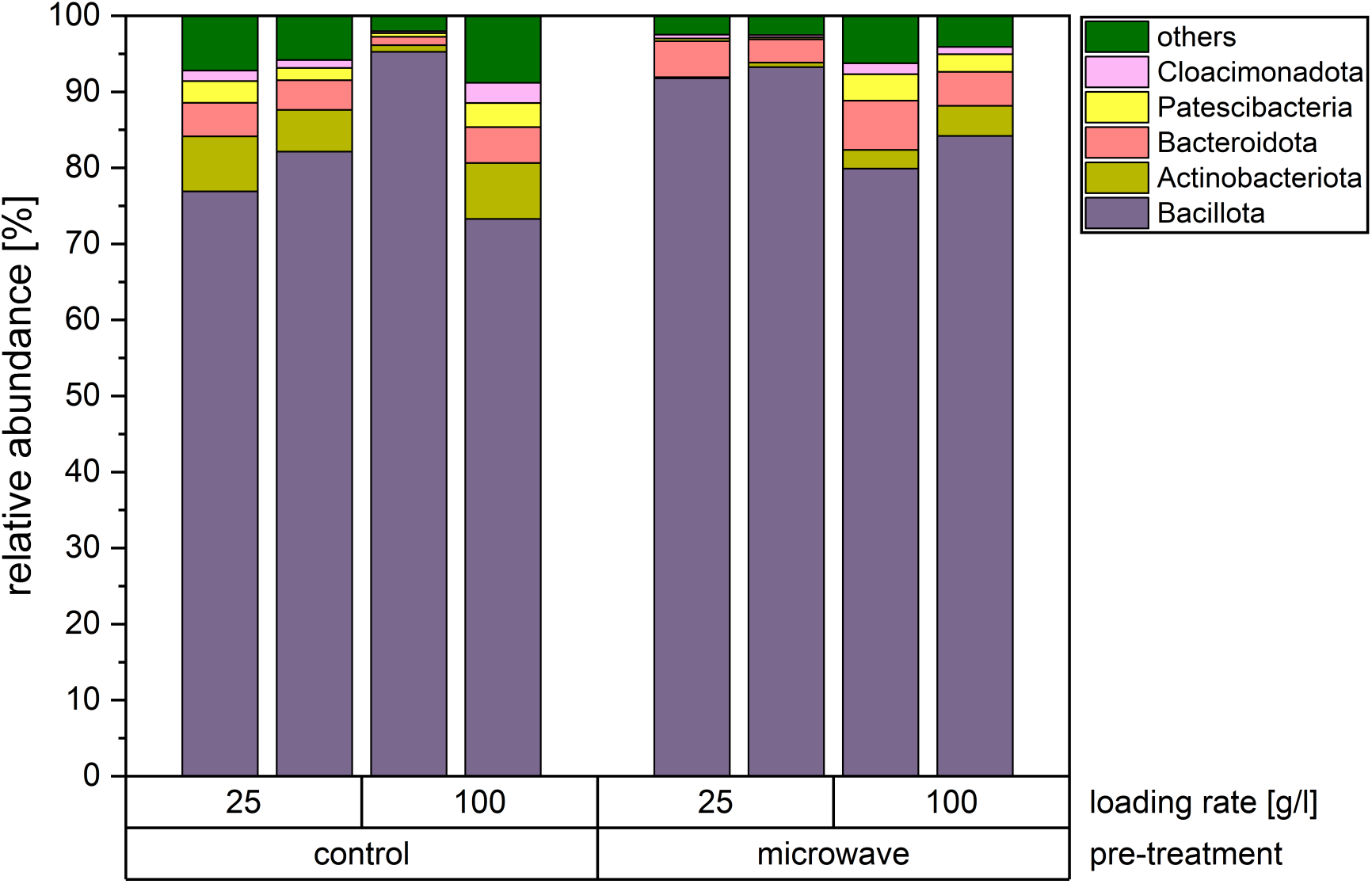
Microbial community on phylum level as relative abundances differentiated according to pre-treatment (loading rates of 25 and 100 g/l). Only the five largest microorganism communities are presented. Other phyla are summarised as “others”.

## Notes

### Competing Interest Statement

The authors have declared no competing interest.

### Summary of Updates

Content has been revised on figures were improved.

